# ASOFormer: a Transformer-based model for predicting antisense oligonucleotide efficacy to support therapeutic candidate prioritization

**DOI:** 10.64898/2026.09.18.752788

**Authors:** Nicholas C. Dove, Yuhao Min, Özkan Iş, Carl Clay, Nilüfer Ertekin-Taner, Daniel P. Wickland, Xue Wang

## Abstract

Antisense oligonucleotides (ASOs) represent a promising therapeutic modality for RNA-based disease mechanisms, but identifying high-efficacy candidates from large sequence spaces remains a major bottleneck in drug development. Here, we present ASOFormer, a novel Transformer-based neural network that predicts the inhibition efficiency of RNase H-mediated ASOs from sequence, predicted secondary structure, and chemical modification features. ASOFormer was trained on a knockdown efficacy dataset spanning over 170, 000 ASO–target pairs compiled largely from published patents. Ablation analysis confirmed that the addition of chemical modification features to the primary sequence drove the largest performance gains, with predicted secondary structure providing complementary benefit when combined with modification information. Further analysis highlighted the importance of the wings of gapmer ASOs and other features. As independent validation, ASOFormer was applied to ASOs targeting seven genes that were not in the training dataset gene list. For four out of the seven genes, the ground truth ASO inhibition efficiency was obtained from a public dataset, whereas the others were measured in-house via qPCR. Compared to two published state-of-the-art methods, ASOFormer achieved the best overall accuracy and prioritized the strongest inhibitors. Crucially, only ASOFormer consistently exceeded random expectation for top-candidate recovery across all seven test genes.

## INTRODUCTION

RNA therapeutics hold great potential to treat human diseases. Antisense oligonucleotides (ASOs), one of the more mature RNA-based therapies, are short sequences of synthetic oligonucleotides that bind to complementary target RNA to modulate gene expression (1, 2). Compared to antibodies and traditional small-molecule therapies, ASOs offer several unique advantages. Because their molecular recognition relies on simple Watson-Crick base pairing, ASOs can in principle be designed to target any gene of interest (1–3). Their straightforward chemical synthesis further accelerates treatment development (4), and ASO-based interventions have now been devised for a wide range of human conditions, including amyotrophic lateral sclerosis (5), retinitis in immunocompromised patients (6), homozygous familial hypercholesterolemia (7), and spinal muscular atrophy (8). In oncology, ASOs are particularly attractive because they can be readily updated to track the evolving mutational landscape of a tumor (3).

ASOs act through one of two principal mechanisms. Gapmer ASOs hybridize to target RNA and form an RNA-DNA heteroduplex and reduce target expression by recruiting RNase H to cleave the bound transcript (1, 2). Steric blocking ASOs, by contrast, bind the target RNA to prevent cellular machinery from accessing a specific region, thereby modulating splicing or translation (1, 2). Because steric blockers must engage defined functional regions of the transcript, they are constrained by where they can bind, whereas gapmers can potentially target a far broader range of sites. This flexibility is also the source of a major challenge: identifying the optimal gapmer for a given target requires searching an enormous space of candidate sequences and chemical configurations, a process that is prohibitively expensive and time-consuming to explore experimentally. Computational tools that can prioritize the most promising candidates are therefore essential for rational ASO design and down-selection.

To address this need, we developed ASOFormer, an artificial intelligence model that predicts the knockdown potential of candidate gapmers against a target RNA. ASOFormer is built on the Transformer architecture, which has driven major advances in natural language processing and is well suited to biological sequence modeling because nucleotides, like words, derive meaning from their context and arrangement (9). The model encodes the primary sequence, predicted secondary structure, sugar modification, and backbone modification of the gapmer, together with the primary sequence and predicted secondary structure of the paired target RNA region, and integrates these features through a hierarchical cross-sequence encoder that captures inter-strand interactions. ASOFormer was trained on over 170, 000 ASO–target pairs from ASO Atlas, a large-scale knockdown efficacy dataset spanning more than 300 genes compiled from licensed patents (10).

Here, we show that ASOFormer accurately ranks ASO knockdown efficacy and reliably prioritizes high-activity candidates. Through systematic ablation, we found that chemical modification features contribute the largest gains in predictive accuracy, with secondary structure providing complementary signal when combined with modification information. Benchmarked against two published state-of-the-art models on seven genes absent from its training data, ASOFormer achieved the best overall rank-order accuracy and top-candidate recall, and was the only model to consistently exceed random expectation for recovering top inhibitors across all test genes. Finally, by applying ASOFormer to an unbiased tiling screen of *KANK2* mRNA and interrogating its learned attention patterns, we demonstrate that the model enriches for efficacious candidates while reducing experimental screening burden, and that its predictions reflect chemically and biologically meaningful sequence features. Together, these results position ASOFormer as a practical tool for therapeutic ASO candidate prioritization.

## RESULTS

### ASOFormer: a Transformer-based model to predict ASO-mediated knockdown

ASOFormer, a Transformer-based neural network, predicts the inhibition efficiency of RNase H-dependent ASOs from sequence and chemical modification features alone. The model accepts two input sequences – the ASO strand and its RNA target binding site – and encodes each using four parallel feature streams and their positional encodings: nucleotide identity, predicted secondary structure, sugar modification, and backbone modification (**Figure 1A**). Sugar and backbone modifications are represented only for the ASO strand; the target strand receives neutral (padding) values for these streams. Feature embeddings at each position are summed prior to positional encoding, allowing the model to share learned representations across combinations rather than learning an entirely separate representation for every possible combination (e.g., the embedding for adenosine is the same regardless of sugar modification).

**Figure 1:**
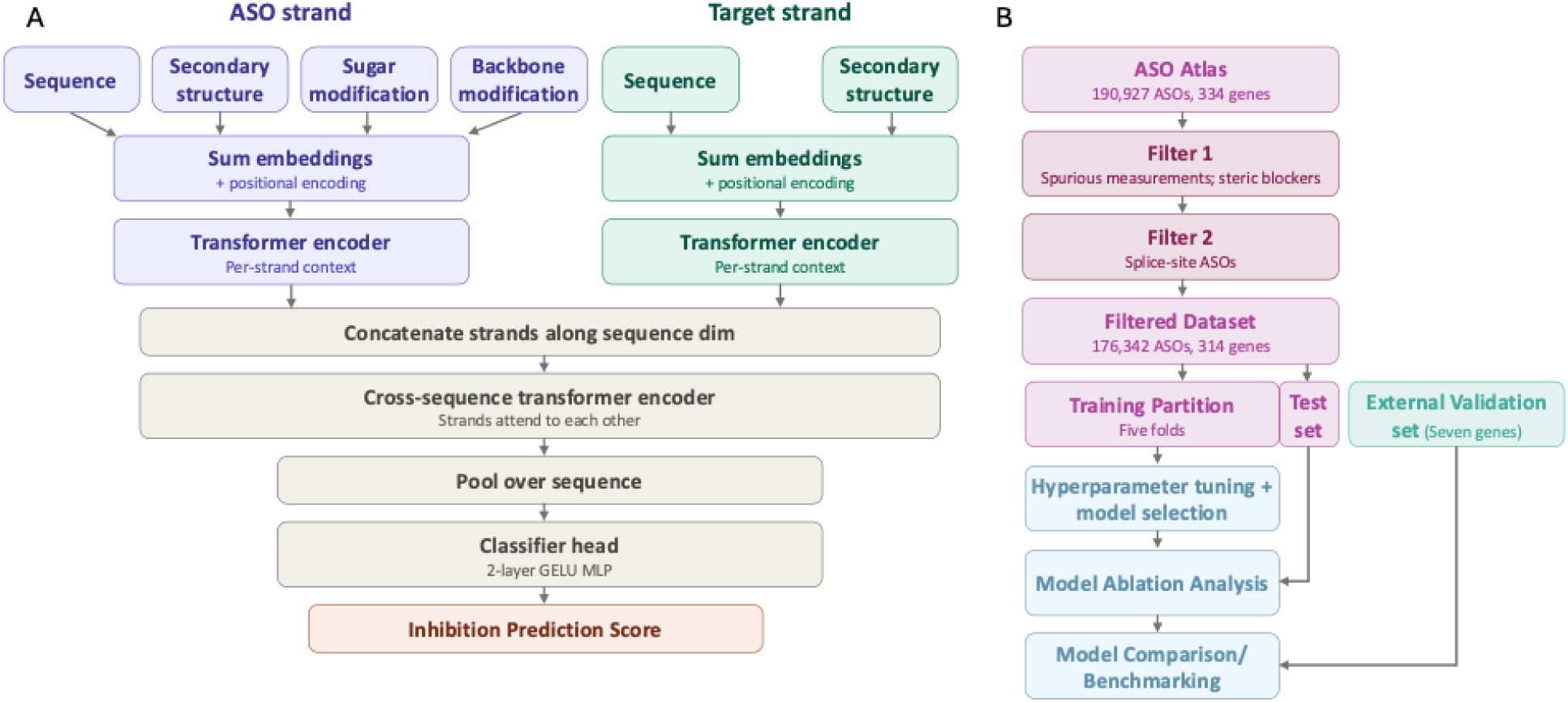
Model architecture and training. (**A**) ASOFormer takes as input the sequence of the ASO and its target binding site, each described by four per-nucleotide feature streams: nucleotide identity, secondary structure, sugar modification, and backbone modification. Chemical modifications are defined only for the ASO strand. Positional encodings are added to the summed embeddings. Each strand is then processed by a dedicated Transformer encoder, after which the two representations are concatenated and passed through a third shared encoder to capture cross-sequence interactions. Padding masks are propagated throughout. The resulting representation is mean-pooled over positions and passed to a prediction head: a two-layer MLP with GELU activation and dropout to produce a scalar inhibition estimate. (**B**) The raw ASO Atlas dataset (190, 927 ASOs targeting 334 genes) was filtered to remove ASOs with inhibition values outside the range of −100% to 100% and those employing a steric blocking mechanism of action (-2, 694 ASOs, 1.4%), followed by exclusion of ASOs spanning splice sites (-11, 891 ASOs, 6.2%), yielding a filtered dataset of approximately 176, 342 ASOs. Prior to model training, this filtered dataset was randomly partitioned into six folds: five used for cross-validation-based model training and hyperparameter selection, and one held out as an internal test set. A second, independent external test set was comprised of in-house and external experimental inhibition measurements across seven genes absent from ASO Atlas.

This compositional approach is parameter efficient and allows the model to generalize across combinations, including those underrepresented in training. After this independent within-strand encoding, representations of the ASO and its target are concatenated and passed through a shared cross-sequence encoder to capture inter-strand interactions, reflecting the complementarity structure of ASO–target binding. The resulting representation is mean-pooled to produce a single fixed-length vector, which is then passed through a two-layer Multilayer Perceptron (MLP) to produce a scalar inhibition prediction.

ASOFormer was trained on 176, 342 RNase H-dependent ASOs drawn from ASO Atlas (10), a large-scale dataset of knockdown efficacy measurements from published patents encompassing 334 unique genes (**Figure 1B**). A held-out random partition of ASO Atlas (one of six equal folds) was withheld prior to any training or hyperparameter tuning. Model selection was performed using 5-fold cross-validation on the training partition (the remaining five of the six equal folds) across a hyperparameter grid (**Table 1**), with Spearman correlation between predicted and observed inhibition as the primary optimization criterion. The final model was trained on all five combined training folds using the optimal configurations. To confirm that training dynamics remained stable with the full training dataset, validation metrics for the held-out partition were monitored throughout the final training run to verify convergence and the absence of overfitting (**Supplementary Figure 1**).

**Table 1:** ASOFormer hyperparameter grid.

| Hyperparameter | Values tested |
| --- | --- |
| Transformer dropout | 0, 0.1, 0.2 |
| Model dimension | 16, 32, 64 |
| Number of layers | 2, 3 |
| Feed-forward dimension | 256, 512 |
| Alpha ($\alpha$) | 0, 1, 2 |

### Secondary structure and chemical modification features improve prediction accuracy

To assess the contribution of each input feature stream, we trained three ablation variants of ASOFormer alongside the full model: primary sequence only, primary sequence with secondary structure, and primary sequence with chemical and backbone modifications. Each variant was trained under the same cross-validation procedure and hyperparameter search as the full model (see Methods: Model Training & Selection).

The sequence-only model achieved a maximum mean cross-fold validation Spearman correlation of 0.393 (validation observed vs. predicted), establishing a baseline that captures the information encoded in nucleotide identity alone (**Figure 2**). Adding predicted secondary structure features improved this figure to 0.402, a modest but consistent gain indicating that the RNA folding context of the ASO and its target contributes signal beyond sequence identity. The larger gain came from adding sugar and backbone modification features in place of secondary structure, which raised the maximum Spearman correlation to 0.462, an improvement of 17.6% over the sequence-only baseline, consistent with the well-established dependence of RNase H activity on ASO chemical modification (11–13). The full model, incorporating all feature streams, achieved the highest maximum validation Spearman correlation of 0.474, outperforming both partial models. That the full model exceeds the sequence-plus-modifications variant (0.474 vs. 0.462) despite secondary structure contributing only modestly on its own (0.393 vs. 0.402) suggests that structural context provides complementary information that is most useful in combination with modification features. Critically, this ordering was consistent across the full range of hyperparameter configurations evaluated, not just at the optimum (**Figure 2**), indicating that the performance advantage of richer feature representations is a robust property of the model rather than an artifact of a particular architectural setting.

**Figure 2:**
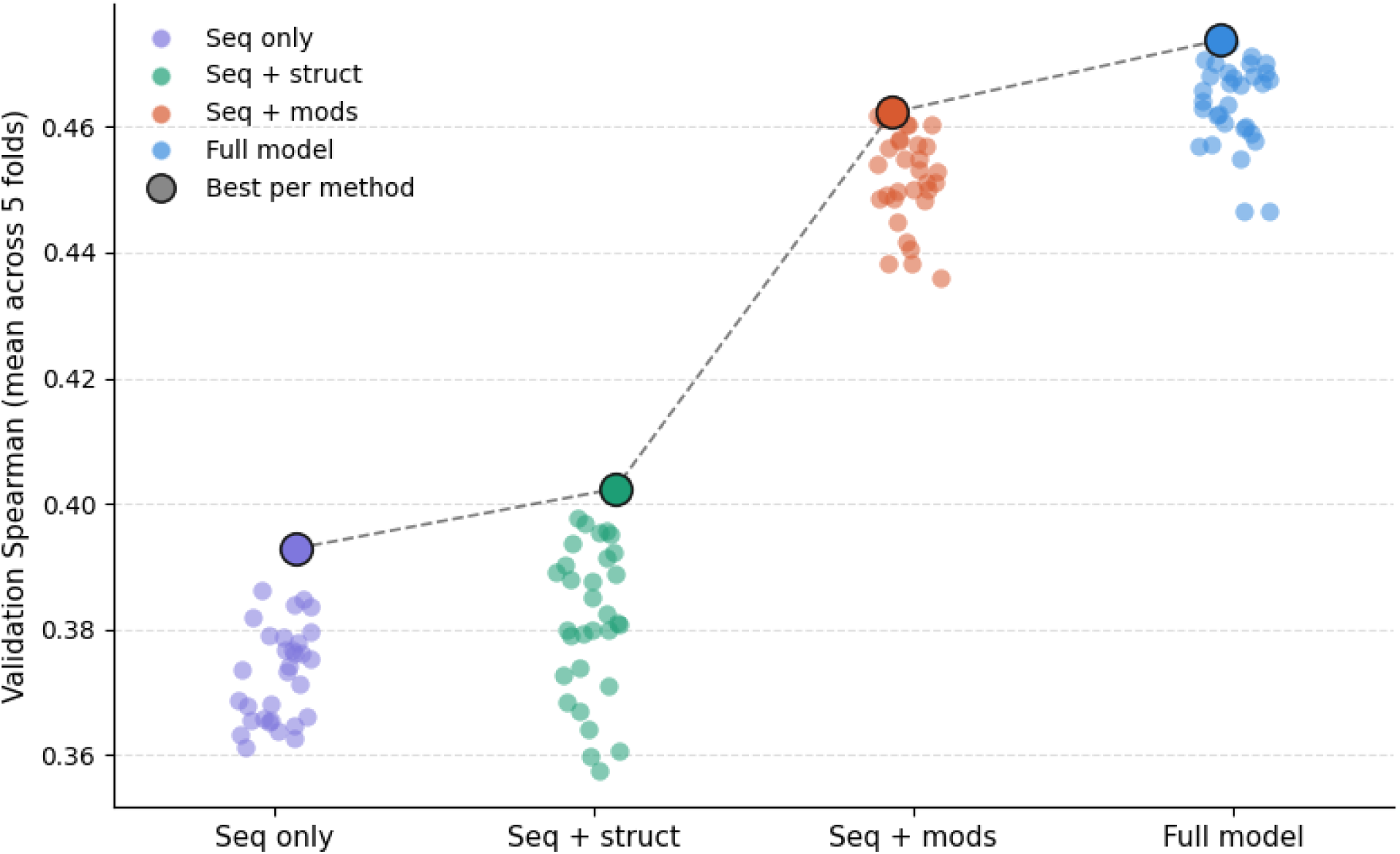
ASOFormer ablation. Mean validation Spearman correlation across five folds for four variants of ASOFormer (sequence-only, sequence and secondary structure, sequence and chemical/backbone modifications, and the full model which includes sequence, secondary structure and chemical/backbone modifications). Each point represents a hyperparameter configuration, with the best configuration (by validation Spearman’s) for each variant highlighted.

### ASOFormer reliably recovers top-activity ASOs across independent gene targets

We next benchmarked the predictive performance of ASOFormer against two previously published ASO efficacy models, ASOptimizer (14) and OligoAI (10), using experimental knockdown data from genes absent from the ASO Atlas training dataset. We nominated 5, 492 candidate ASOs from seven such genes and compared the *in silico* predicted knockdown under each model to the experimentally measured knockdown data. The latter we obtained from both in-house experiments (*DDR2*, *IQGAP3*, and *KANK2*, **Supplementary Table 1**, **Supplementary Figure 2**) and OligoGym (*ANGPTL2*, *SNHG14*, *HTRA1*, and *MYH7*) (15). *DDR2*, *IQGAP3* and *KANK2* were included in all three-way model comparisons; however, the four genes sourced from OligoGym were not used to benchmark ASOptimizer because its training dataset (Hwang et al. (14)) likely included them. All model weights were used as released, without retraining or fine-tuning. Performance was evaluated using two complementary metrics: Spearman correlation, which measures overall rank-order accuracy, and top-10% recall, which measures the proportion of the true top-decile inhibitors recovered when selecting the top 10% of ASOs by predicted score. The latter metric reflects practical utility for candidate prioritization, where failing to identify high-activity ASOs impairs therapeutic development.

Across these test genes, ASOFormer achieved a mean Spearman correlation of 0.268 (range: −0.029 to 0.534) between observed and predicted inhibition, compared to 0.223 (range: −0.125 to 0.508) for OligoAI and 0.055 (range: −0.442 to 0.493) for ASOptimizer, representing a 20.2% improvement over OligoAI and xx% improvement over ASOptimizer in mean rank-order accuracy (**Figure 3A**). While these correlations are modest in absolute terms, rank-order accuracy across the full range of inhibition values is a demanding criterion that weights performance on both inactive and highly active ASOs equally. Instead, a metric more directly relevant to practical screening is top-10% recall, which quantifies the model’s ability to reliably identify the small subset of ASOs with the highest activity. The differences between models based on this measure were stark (**Figure 3B**). A model with no predictive power would achieve a recall of 0.1 by chance. ASOFormer met or exceeded this baseline across every test gene, with a mean recall of 0.259 (range: 0.108–0.667), demonstrating consistent above-chance identification of high-activity ASOs regardless of target. OligoAI achieved a nominally above-baseline mean recall of 0.138, but this value was driven entirely by strong performance on a single gene (*IQGAP3*, recall: 0.667); on the remaining genes, OligoAI fell below random expectation. ASOptimizer performed near or below random chance on most targets, achieving a mean recall of just 0.037 and reaching approximately baseline on only one gene (*KANK2*, recall: 0.111). Spearman correlations did not differ significantly between ASOFormer and OligoAI (pairwise Wilcoxon, *p* = 0.688; comparisons with ASOptimizer were not included due to too few test genes).

**Figure 3:**
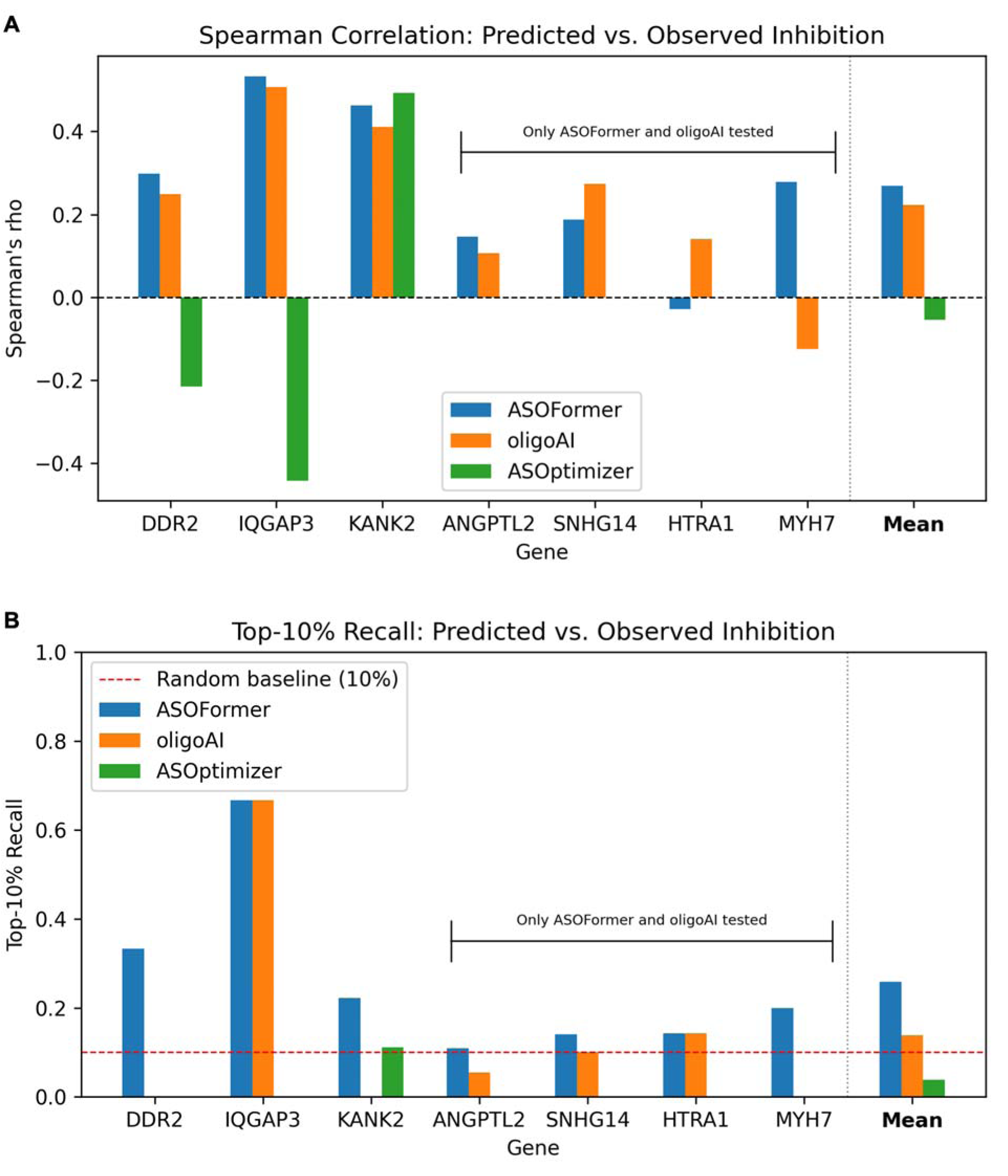
Model Validation. Spearman correlation (A) and top-10% recall (B) of ASOFormer prediction accuracy compared with oligoAI and ASOptimizer using knockdown data from seven genes not represented in the training dataset (ASO Atlas). Genes *ANGPTL2*, *SNHG14*, *HTRA1*, and *MYH7* were used in the training of ASOptimizer and were not used in this evaluation. ASOFormer achieved the highest mean Spearman correlation (0.268) and top-10% recall (0.259) across all test genes, consistently exceeding random expectation for identifying high-activity ASOs.

### ASOFormer cross-encoder attention patterns are consistent across three target genes

To assess whether ASOFormer’s cross-encoder attention weights reflect interpretable sequence features, we extracted per-position attention weights from the cross-sequence encoder across all internally screened ASOs (*DDR2*, *IQGAP3*, *KANK2*). Attention weights in the cross-sequence encoder reflect the degree to which the model references each target position when constructing its representation of each ASO position, forming a matrix of size ASO length × target length (20×20 for each ASO-target pair). These weights provide a window into which sequence features the model considers contextually relevant during inter-strand reasoning, though they do not directly quantify contribution to the inhibition prediction. We summarized these matrices by summing attention over one strand dimension to obtain per-position attention profiles for each strand and assessed how attention varied across ASO chemical modification regions, prediction outcome groups, and target nucleotide identity (**Supplementary Figure 3**).

We summed the attention weights for each ASO base across the target strand. The first and last five positions of the ASO sequence, representing the 5′ and 3′ MOE wings, received significantly higher attention than the central DNA gap across all three gene targets (Wilcoxon signed-rank, Bonferroni-corrected *p_adj_*< 0.002 in *DDR2* and *IQGAP3*, *p_adj_* < 0.001 in *KANK2*); the two wings did not differ from each other in any gene (**Figure 4A**). These observations suggest that the model preferentially encodes modification context at the wing–gap boundary during inter-strand reasoning, consistent with the known role of MOE flanks in RNase H recruitment and heteroduplex stability (11–13). We also summed the attention weights for each target strand base across the ASO strand. G residues in the target sequence attracted substantially higher attention weights than all other nucleotides in all three genes (Kruskal-Wallis, all *p* < 0.001; **Figure 4B**).

**Figure 4:**
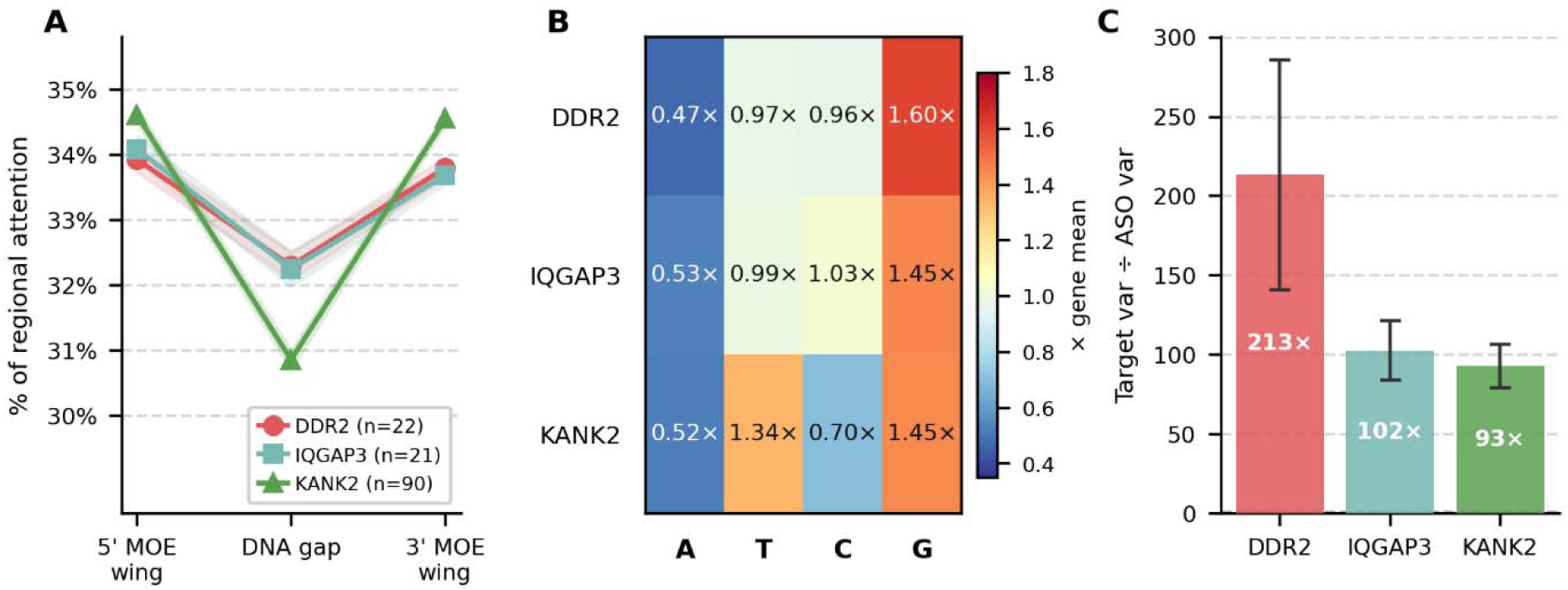
Cross-encoder attention analysis across *DDR2*, *IQGAP3*, and *KANK2* ASOs. (**A**) Attention from each ASO position to the target RNA, summed over all target positions and expressed as a percentage of the total attention distributed across the three gapmer regions. Lines show the mean across ASOs; shaded bands, ±SEM. (**B**) Mean attention received by each target nucleotide identity, normalized row-wise to the gene mean (values indicate fold-change relative to that gene’s mean attention per nucleotide). (**C**) Positional specificity of cross-encoder attention on the target strand relative to the ASO strand, quantified as the per-ASO ratio of positional variance (target ÷ ASO). Error bars, ±SEM. Numbers inside bars indicate the mean fold-change. MOE, 2′-*O*-methoxyethyl; SEM, standard error of the mean.

The relative ordering of T and C was not statistically distinguishable in any gene (all pairwise *p_adj_* > 0.05), yielding a consistent hierarchy of G > T/C > A. This ordering is notable given the well-documented decline in efficacy of ASOs at high GC content (16), suggesting the model has learned to weight G-rich target positions as informative features, potentially as a proxy for the inhibitory effects of excessive heteroduplex stability. The mean positional variance of target-side attention profiles was approximately 60- to 80-fold higher than that of ASO-side profiles across all three gene targets (Wilcoxon signed-rank, all *p* < 0.001; **Figure 4C**), indicating that the model strongly differentiates among target positions but treats ASO positions relatively uniformly once modification context is accounted for. This suggests that inter-strand reasoning is primarily organized around target-site features rather than ASO-intrinsic sequence identity. Together, these results demonstrate that ASOFormer’s inter-strand attention captures chemically and biologically meaningful sequence features that are reproducible across three independent gene targets, lending greater confidence to their mechanistic interpretation.

### ASOFormer prioritizes top-performing ASOs from an unbiased tiling screen of *KANK2*

To demonstrate the practical utility of ASOFormer in a prospective screening workflow, we applied the model to retroactively prioritize ASOs targeting randomly tiled regions of *KANK2* exons. Rather than relying on heuristic sequence rules or expert intuition, we designed an unbiased tiling library to achieve comprehensive coverage of the *KANK2* transcript (ENST00000586659.6). Specifically, we selected up to 10 non-overlapping 20-nucleotide ASO binding sites per exon across all 13 annotated exons, dividing each exon into equal-length segments and sampling one candidate site per segment, yielding a final panel of 90 ASOs spanning the exonic sequence space (**Supplementary Table 1**). This tiling strategy ensured that no region of the transcript was systematically under- or over-represented, providing a ground truth reference for evaluating model-guided prioritization.

Each of the 90 tiled ASOs was scored by ASOFormer and ranked by predicted inhibition percentage. Consistent with the recall performance described above (**Figure 3B**), we defined top performing ASOs as those exceeding the 90th percentile of observed inhibition, and model-predicted positives analogously as those exceeding the 90th percentile of ASOFormer scores.

ASOFormer recovered two of the nine top-performing ASOs among its top nine predictions, corresponding to a recall of 0.222 versus a random expectation of 0.1 (2/9 recovered vs. 9/90 expected by chance; 2.2-fold enrichment). This observation suggests that restricting experimental validation to the top-decile of model-ranked candidates (10% of the full library) could recover a substantial proportion of high-activity ASOs while reducing screening burden by approximately 90%. Notably, the remaining high-confidence ASOFormer predictions (false positives by the strict 90th percentile criterion) were nonetheless among the higher-performing ASOs experimentally, suggesting the model’s top predictions are broadly enriched for efficacious candidates (**Figure 5A**). ASOs with low observed inhibition were largely confined to lower predicted scores and would not be advanced in a model-guided workflow regardless.

**Figure 5:**
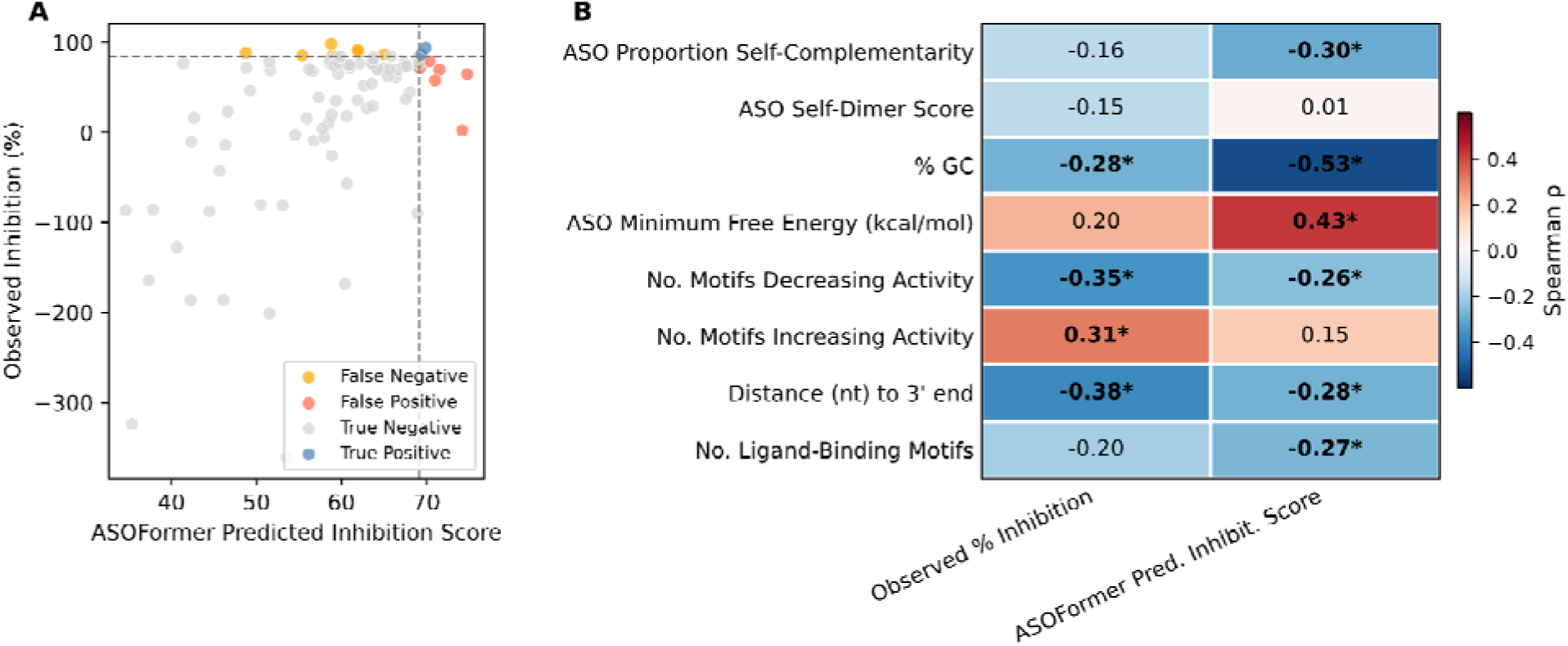
Investigation of the *KANK2* unbiased tiling library. (**A**) Observed versus predicted values with points (ASOs) colored by whether they are represented in the top 10% of both predicted and observed values (True Positive), top 10% of predicted values but outside the top 10% of observed values (False Positive), top 10% of observed values but outside of the top 10% of predicted values (False Negative), or represented in the top 10% of neither predicted nor observed values (True Negative). (**B**) Correlations (Spearman’s rho) between *KANK2* ASO characteristics and observed percent inhibition or ASOFormer predicted inhibition score. Significant correlations (*p_adj_* < 0.05) are indicated by an asterisk (*).

To further characterize the features driving ASOFormer predictions and their relationship to observed efficacy, we compiled a panel of ASO physicochemical and sequence characteristics and computed their Spearman correlations with both observed percent inhibition and predicted inhibition score across the *KANK2* tiling library (**Figure 5B**). Several features were significantly correlated with both metrics, suggesting they capture biologically meaningful determinants of ASO activity. Among sequence composition features, GC percentage showed the strongest negative associations with both observed inhibition (rho = −0.28, *p_adj_* < 0.013) and predicted score (rho = −0.53, *p_adj_* < 0.001), a result consistent with the well-documented decline in efficacy of ASOs at high GC content (16). Three measures of ASO self-structure showed no significant correlation with observed inhibition percentage: 1) proportion self-complementarity (the fraction of bases involved in intramolecular binding structures in the predicted secondary structure), 2) self-dimer score (17), and 3) mfold-derived minimum free energy (18). This lack of association suggests that self-structure may not impact efficacy for this particular target. However, two of these self-structure measures reached significant association with the ASOFormer predicted score (proportion self-complementarity: rho = −0.30; minimum free energy: rho = 0.43, both *p_adj_* < 0.05), suggesting the model has learned to weight self-structure based on patterns in its broader training data. This discrepancy may also reflect gene-level dependence of self-structure effects across targets, where self-structure is less predictive for *KANK2* ASOs, though further validation across additional transcripts is required to confirm this possibility.

In addition to self-structure, binding site start position was negatively correlated with both observed and predicted inhibition (rho = −0.38 and −0.28, respectively, both *p_adj_* < 0.05, **Figure 5B**). For *KANK2*, which is encoded on the negative strand, lower genomic coordinates correspond to the 3′ end of the transcript; therefore, the observed negative correlation reflects modestly greater activity at 3′-proximal target sites in this library. The number of activity-decreasing motifs (compiled from (19, 20); see Methods) was negatively associated with both observed and predicted knockdown (rho = −0.35 and −0.26, respectively, both *p_adj_* < 0.05), while the number of activity-increasing motifs showed a positive association, consistent with the sequence-motif rules from which these features were derived. The broad concordance between feature correlations for observed and predicted values further implies that ASOFormer learned to weight these properties in a manner consistent with experimental outcomes, rather than capturing spurious patterns. Notably, while several individual ASO characteristics reached statistical significance, none achieved a Spearman correlation with observed inhibition as strong as ASOFormer’s overall ranking (rho = 0.463, **Figure 3A**), suggesting that the model’s predictive power derives from integrating patterns across multiple sequence and structural features simultaneously rather than from any single dominant attribute.

## DISCUSSION

Here we present ASOFormer, a Transformer-based model that predicts the inhibition efficiency of RNase H–dependent ASOs from sequence and chemical modification features alone. Trained on more than 170, 000 ASO–target pairs from ASO Atlas, ASOFormer achieved the best overall rank-order accuracy and the most reliable recovery of top-activity candidates when benchmarked against two published state-of-the-art models on seven genes absent from its training data. Critically, it was the only method to consistently exceed random expectation for top-candidate recovery across every test gene, as indicated by its superior top-10% recall compared to the other two models. Because the central bottleneck in gapmer development is the cost and time of experimentally searching an enormous candidate space, a model that more reliably enriches for the most active candidates addresses the problem in the terms that matter for drug screening.

A consistent theme across our analyses is that ASOFormer’s predictive power is anchored in ASO chemistry, which the model appears to have learned in a biochemically faithful way. In ablation, chemical modification features drove the largest single gain in accuracy, while predicted secondary structure contributed only modestly alone and was most useful in combination with modification information. This ordering mirrors decades of biochemical work showing that RNase H recruitment and heteroduplex processing depend heavily on the chemistry and placement of sugar and backbone modifications rather than on nucleotide identity in isolation (11, 21). Strikingly, the model’s internal attention patterns recapitulate the same biology without it being explicitly modeled. The cross-encoder directed disproportionate attention to the 5′ and 3′ MOE wings relative to the central gap, concentrating on the wing–gap boundary that governs heteroduplex stability and RNase H engagement (11, 21). The accuracy gain attributable to chemistry features and the attention pattern centered on the chemically modified wings arise independently yet point to the same conclusion. That convergence provides greater confidence that ASOFormer captures a genuine biophysical determinant of activity rather than a dataset artifact, consistent with reports that chemically optimized designs can substantially outperform conventional gapmers of identical sequence (14).

This learned biochemistry extends to the target strand as well. The model weighted target G residues most heavily (a G > T/C > A hierarchy) and structured its attention far more strongly along the target than the ASO strand, indicating that its inter-strand reasoning is organized primarily around target-site context. This observation aligns with independent evidence that excessive GC-driven heteroduplex stability can impair efficacy (16) and that human RNase H1 exhibits reproducible nucleotide-level cleavage preferences correlated with gapmer activity (22). We caution that attention weights indicate where the model attends, not the magnitude or direction of a feature’s contribution, so these patterns are best read as hypotheses about the model’s learned representations rather than causal claims; their reproducibility across three independent gene targets nonetheless lends them interpretive weight.

The second theme is that ASOFormer’s utility derives from integrating many individually weak determinants rather than from any single dominant rule, making it valuable in a screening workflow. In the *KANK2* feature analysis, GC content, self-structure measures, binding-site position, and curated activity-associated motifs each correlated with predicted score in directions concordant with their known relationships to efficacy (16, 19, 20), yet none individually approached the rank-order accuracy of the model’s integrated prediction. This is the regime in which learned multi-feature models are expected to outperform rule-based design (23). The unbiased *KANK2* tiling screen shows the practical consequence: applied to a position-balanced library built to remove the heuristic biases that normally shape candidate panels, ASOFormer enriched for top-decile inhibitors 2.2-fold over chance, with its remaining high-confidence calls clustering among the higher-performing ASOs even when they fell outside the strict cutoff. The implication for deployment is direct: restricting experimental validation to the top decile of model-ranked candidates could recover a substantial share of the strongest inhibitors while cutting screening burden by roughly 90%. This is the same property that distinguishes ASOFormer in benchmarking, where it was the only model to reliably exceed random expectation for top-candidate recovery across every test gene.

Several limitations of our study warrant consideration. The absolute correlations achieved by ASOFormer and all comparator models were moderate (mean Spearman ∼0.27), reflecting both the intrinsic noise of pooled, cross-study knockdown measurements and the difficulty of generalizing to unseen genes. Our external benchmark comprised only seven genes, which limited statistical power; the difference in top-10% recall between ASOFormer and OligoAI was striking in practical terms, but the paired Spearman comparison did not reach significance, and the three-way comparison with ASOptimizer was constrained because four OligoGym genes appeared in its training data. The training data themselves, derived from patent-extracted measurements in ASO Atlas (10), are heavily concentrated near median inhibition with sparse tails, which limits our ability to predict actual inhibition percentages. We partially addressed this limitation through tail-weighted loss (see Methods), but resolution at the extremes of the activity range nevertheless remained coarse. Finally, the model predicts only *in vitro* knockdown efficacy and does not address the equally consequential dimensions of off-target activity, hepatotoxicity, and *in vivo* delivery that ultimately gate therapeutic development (24, 25).

These limitations clearly point to future extensions of this work. Expanding training data to include diverse chemistries and a richer representation of the activity tails, jointly modeling efficacy alongside toxicity and off-target liability, and prospectively validating model-prioritized candidates *in vivo* would move ASOFormer closer to an end-to-end design tool. The capacity to generalize to genes never seen in training (the property most directly relevant to deployment) was itself improved here by random (vs. gene-stratified) data splitting (**Supplemental Note 1 Supplementary Figure 4**), an empirical result suggesting that data scale and cross-gene diversity, rather than enforced gene holdout, currently provide better generalizability. As ASO efficacy datasets continue to grow (10), we anticipate that this advantage will widen. Together, our results position ASOFormer as a practical tool for prioritizing therapeutic ASO candidates: it reliably enriches for high-activity gapmers, reduces experimental screening burden, and bases its predictions on chemically and biologically interpretable features.

## METHODS

### Model Architecture

ASOFormer is a Transformer-based neural network written in PyTorch (26) and designed to predict the inhibition efficiency of antisense oligonucleotides (ASOs) (26). The model accepts two input sequences – the ASO strand and its RNA target strand – and processes them through a hierarchical encoding and attention framework before producing a scalar inhibition prediction.

Each nucleotide position is described by four parallel feature streams: nucleotide identity, predicted secondary structure, sugar modification, and backbone modification. Each stream has its own independent embedding table, with index 0 reserved as a padding token across all vocabularies. The nucleotide vocabulary encodes C, A, G, T, and N (five tokens); the secondary structure stream encodes unpaired positions and opening/closing base-pair brackets in dot-bracket notation (three tokens), with secondary structures predicted using mfold from the ViennaRNA package; the sugar modifications include MOE, DNA, cEt, OMe, and 2’-F (five tokens); and the backbone modification stream encodes phosphodiester (PO) and phosphorothioate (PS) linkages (two tokens).

These four embedding vectors at each position are summed prior to downstream processing. This additive composition is not novel to our architecture; it follows the same strategy used by BERT, which sums token, segment, and positional embeddings into a single input representation (27). Adopting this well-established design lets the model learn independent representations for nucleotide identity and for each chemical modification. It therefore avoids having to learn a separate representation from scratch for every possible nucleotide × modification combination. The target strand, which carries no chemical modifications, has its sugar and backbone arrays set to zero (the padding index), so the model receives a clean signal on that strand without spurious modification tokens. Fixed sinusoidal positional encodings (sine and cosine functions of varying frequency) are added element-wise to the summed input embedding at each position, giving the otherwise order-agnostic self-attention mechanism information about token order.

The encoded ASO and target strands are first processed independently through separate Transformer encoder blocks, allowing each strand to develop contextual representations within itself. The two independently encoded representations are then concatenated along the sequence dimension and passed through a third, cross-sequence Transformer encoder. All transformer encoders use two attention heads. This hierarchical design enables the model to build intra-strand context before attending to inter-strand relationships, which is well-suited to the complementarity structure of ASO–target binding. Padding masks derived from the nucleotide index arrays are propagated through all three encoders so that padded positions do not contribute to attention computations.

Following the cross-sequence encoder, the sequence representation is reduced to a fixed-length vector via mean pooling over non-padded positions. The pooled representation is then passed through a two-layer Multilayer Perceptron (MLP) regression head with Gaussian Error Linear Unit (GELU) activation and dropout to produce a scalar inhibition prediction.

### Training Data

ASOFormer was trained on the ASO Atlas dataset, a collection of 190, 927 RNase H-mediated ASO sequences targeting 334 unique genes, with corresponding knockdown efficacy measurements extracted from published patents (10).

Prior to model training, the dataset was filtered to remove likely spurious measurements. Specifically, we removed ASOs with inhibition percentages outside the range of −100% to 100%. ASOs employing a steric blocking mechanism of action were also excluded, as these operate via distinct biophysical principles that differ from the RNase H-dependent cleavage activity represented by the majority of the dataset, and their inclusion would complicate the regression objective. Together, these filters removed 2, 694 ASOs (1.4% of the dataset).

Secondary structure features were computed as described in the Model Architecture section. ASO secondary structure was predicted using ViennaRNA’s mFold algorithm under default parameters (18). Binding site secondary structure was derived from the full unspliced gene sequence, computed using ViennaRNA’s lFold algorithm with the --backtrack-global flag enabled and all other parameters set to default, then clipped to the binding site coordinates provided by ASO Atlas. ASOs spanning splice sites could not be assigned a valid binding site structure under this approach and were excluded, both for this technical reason and because splice-switching ASOs represent a mechanistically distinct class beyond the scope of this study. This step removed an additional 11, 891 ASOs (6.2% of the complete dataset). Secondary structure strings were further validated prior to training to confirm that they contained only valid dot-bracket characters and that modification arrays were of consistent length with their corresponding sequences. After all filtering, approximately 176, 342 ASOs were retained for model training and evaluation.

Two independent test sets were established prior to any model training or hyperparameter tuning and were reserved exclusively for final performance evaluation. The first consisted of a held-out random portion of the ASO Atlas dataset. The second comprised experimental inhibition measurements generated in house (described in Experimental Methods). Neither test set was used at any stage of model selection.

### Model Training & Selection

Models were trained on an NVDIA H100 cluster and selected using 5-fold cross-validation. At each fold, one partition was held out as the validation set and the remainder was used for training. The ASO Atlas training partition was defined by random split. Gene- and patent-level stratification were also evaluated; however, both stratified models demonstrated inferior generalization to the external test set, suggesting that at the scale of ASO Atlas, random splitting preserves sufficient cross-gene diversity to support generalizable training without meaningful leakage inflation. The random-split model was therefore selected for all subsequent analyses (**Supplementary Note 1, Supplementary Figure 4**).

Because the ASO Atlas dataset is heavily concentrated near median inhibition values with the tails of the distribution substantially underrepresented (**Supplementary Figure 5**), a tail-weighted loss function was used to improve prediction accuracy across the full activity range. Each training example was assigned a sample weight proportional to its distance from the median inhibition value of the training set:

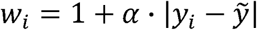

where y*_i_*, is the inhibition proportion for sample *i*, ỹ is the median inhibition proportion across the training set, and α is a scaling factor controlling the magnitude of the tail-weighting effect, with larger values placing greater emphasis on ASOs with extreme inhibition values. Weights were normalized to have unit mean within each batch before computing the loss. Models were optimized using Adaptive Moment Estimation (Adam) with a learning rate of 1×10. Gradient norms were clipped to 0.5 to stabilize training.

Validation performance was assessed every 10 epochs using Spearman correlation between predicted and observed inhibition on the held-out fold. Training was halted if validation Spearman correlation failed to improve by more than 0.0001 over five consecutive validation checks (50 epochs without meaningful improvement). The model checkpoint with the highest validation Spearman correlation was retained.

Five hyperparameters were tuned by training a separate model for each combination across all five folds (**Table 1**). The selected configuration was that which achieved the highest mean validation Spearman correlation averaged across all five folds, with mean validation RMSE used as a tiebreaker.

The final models were trained on all five combined training folds using the optimal configurations from **Table 1**. To confirm that training dynamics remained stable with the full training dataset, validation metrics were monitored throughout the final training run to verify convergence and the absence of overfitting (**Supplementary Figure 1**). A similar early-stopping heuristic was also implemented.

### Model Ablation

To assess the contribution of each input feature stream, three ablation variants of ASOFormer were trained and evaluated alongside the full model:

- **Sequence only**: secondary structure and chemical modification features (sugar and backbone) excluded
- **Sequence + secondary structure**: chemical modification features (sugar and backbone) excluded
- **Sequence + chemical modifications**: secondary structure excluded

Each ablation variant was trained under the same procedure described in Model Training & Selection, using 5-fold cross-validation and a reduced hyperparameter grid informed by prior experimentation. The best-performing configuration for each variant was selected using the same criterion as the full model: highest mean validation Spearman correlation across all five folds, with mean validation RMSE as a tiebreaker.

### Experimental Methods

#### ASO Synthesis

We synthesized 133 ASOs targeting three genes: *DDR2*, *KANK2*, and *IQGAP3*. The sequences of all ASOs used in this study are provided in **Supplementary Table 1**. All ASOs were 20 nucleotides in length with a full phosphorothioate backbone. The five terminal nucleotides at both the 5′ and 3′ ends were modified with 2′-*O*-methoxyethyl (2′-MOE) groups, flanking a central 10-nucleotide DNA gap region (a gapmer design). ASOs were synthesized by Integrated DNA Technologies (IDT), reconstituted in TE buffer to a stock concentration of 100 µM, and stored at −20°C until use.

#### KANK2 Tiling Library Design

To enable an unbiased, position-independent evaluation of ASOFormer’s prioritization performance, we designed a systematic tiling library targeting the *KANK2* transcript (ENST00000586659.6). For each of the 13 annotated exons, we selected up to 10 non-overlapping 20-nucleotide ASO binding sites by dividing each exon into equal-length segments and sampling one candidate site per segment. This strategy yielded a final panel of 90 ASOs spanning the full exonic sequence space, with no region of the transcript systematically under- or over-represented. All 90 ASOs followed the same gapmer chemistry described above (full phosphorothioate backbone; 2′-MOE modifications at the five terminal nucleotides on both ends flanking a 10-nucleotide DNA gap). Binding site coordinates and ASO sequences for the complete tiling library are provided in **Supplementary Table 1**.

#### Cell Culture

Human neuroglioma H4 cells were maintained at 37°C under 5% CO in Opti-MEM (Thermo Fisher Scientific, cat. no. 51985034) supplemented with 10% fetal bovine serum (Thermo Fisher Scientific, cat. no. A5256701). Cells were passaged at 70–90% confluency using Trypsin (Thermo Fisher Scientific, cat. no. 25200072).

#### ASO Transfection

H4 cells were seeded onto tissue culture-treated well plates at a density of 30, 000 cells/cm² and allowed to adhere for 24 hours prior to transfection. ASOs were delivered at a final concentration of 100 nM using Lipofectamine 3000 Transfection Reagent (Thermo Fisher Scientific, cat. no. L3000115) according to the manufacturer’s protocol with minor modifications. Briefly, equal volumes of 2 µM ASO working solution and 6% (v/v) Lipofectamine 3000 working solution, each prepared in Opti-MEM, were combined, mixed gently, and incubated at room temperature for 15 minutes before being added dropwise to cells. Each ASO was transfected and tested in H4 cells for three times independently.

#### RNA Extraction and RT-qPCR

Total RNA was extracted using the RNeasy Mini Kit or RNeasy 96 Kit (Qiagen, cat. nos. 74104 and 74181, respectively) following the manufacturer’s instructions. For samples designated for RNA sequencing, genomic DNA was eliminated via on-column DNase digestion using the RNase-Free DNase Set (Qiagen, cat. no. 79254). For extractions performed with the RNeasy 96 Kit, all centrifugation steps were conducted at a reduced speed of 3, 486 × *g* for 7 minutes, and an additional RPE wash step was included to improve RNA purity. RNA concentration and purity were assessed using a NanoDrop 2000c spectrophotometer (Thermo Fisher Scientific).

Complementary DNA (cDNA) was synthesized from 20-2000 ng RNA per 20 uL reaction using the High-Capacity cDNA Reverse Transcription Kit (Thermo Fisher Scientific, cat. no. 4368813) per the manufacturer’s instructions. Quantitative PCR (qPCR) was performed on a QuantStudio 7 Flex Real-Time PCR System (Thermo Fisher Scientific) using TaqMan Fast Advanced Master Mix (Thermo Fisher Scientific, cat. no. 4444557) with pre-designed TaqMan gene expression assays (**Supplementary Table 2**). The qPCR reactions were performed in duplex, with probes for both the genes of interest (*DDR2*, *IQGAP3*, or *KANK2*) and the housekeeping gene (*ACTB*). The reactions were run in technical triplicates, where the mean CT values from the technical replicates were used for quantification. Relative gene expression downregulation was calculated using the delta-delta CT methods (https://www.nature.com/articles/nprot.2008.73) using ACTB as the housekeeping gene.

### Model Comparison

ASOFormer was benchmarked against two existing ASO efficacy prediction models: ASOptimizer (14) and OligoAI (10). Trained model weights were obtained directly from the respective public repositories (ASOptimizer: github.com/Spidercores/ASOptimizer; OligoAI: huggingface.co/barneyhill/OligoAI) without any retraining or fine-tuning.

### Statistical Analysis

All statistical analyses were performed in Python (v3.10). Data visualization was generated using Matplotlib (v3.8.4) (28) and Seaborn (v0.13.2) (29). The ASOFormer model was implemented in PyTorch (v2.3) (26).

Rank-order predictive performance was assessed using Spearman’s rank correlation coefficient (rho), computed between predicted and observed ASO inhibition values. Spearman correlation was selected over Pearson correlation given the ordinal nature of model ranking utility and the need to limit sensitivity to outliers in inhibition measurements. Spearman correlations were computed for each gene independently, and mean performance across genes is reported alongside the observed range.

Top-10% recall was calculated as the proportion of ASOs with observed inhibition values in the top decile that were also ranked in the top decile by predicted score. This metric was chosen as the primary measure of practical screening utility, as it directly quantifies a model’s ability to enrich for high-activity candidates. Pairwise comparisons of top-10% recall between ASOFormer and OligoAI across the seven test genes were performed using the Wilcoxon signed-rank test, a non-parametric test appropriate for paired observations from a small sample. The same test was applied to Spearman correlations.

To characterize the relationship between ASO properties and knockdown efficacy, we assessed a panel of physicochemical and sequence features for all 90 ASOs in the *KANK2* tiling library. GC percentage was calculated directly from each ASO sequence. ASO secondary structure was predicted using ViennaRNA’s mfold algorithm under default parameters, from which we derived minimum free energy (kcal/mol) and proportion self-complementarity, defined as the fraction of nucleotide positions predicted to participate in intramolecular base-pair interactions in the minimum free energy structure. Self-dimer score was computed as described in (17). Binding site start position was extracted from genomic coordinates (GRCh38). The number of sequence motifs associated with decreased or increased ASO activity was calculated by scanning each ASO sequence against curated motif lists compiled from (19, 20).

Spearman rank correlations were computed between each feature and both observed percent inhibition and ASOFormer predicted inhibition score across the 90 *KANK2* tiling ASOs. Multiple-testing correction was applied using the Benjamini–Hochberg false discovery rate (FDR) procedure across all features within each outcome variable. Corrected p-values (*p_adj_*) are reported, and correlations with *p_adj_* < 0.05 were considered statistically significant.

## ACKNOWLEDGMENT

The authors gratefully acknowledge the funding support from the Mayo Clinic RNA Therapeutics Discovery and Translation Program. We also thank Drs. Yan W. Asmann, Margot A. Cousin, Tsuneya Ikezu, and Tushar C. Patel for their valuable input.

## DATA & SOFTWARE AVAILABILITY

The ASOFormer model, training scripts, model weights, and analysis notebooks that were used to generate figures in this manuscript can be found at https://github.com/nicholascdove/asoformer. ASO Atlas was downloaded from https://github.com/barneyhill/aso_atlas, and OligoGym was downloaded from https://github.com/Roche/OligoGym. For comparison with OligoAI and ASOptimizer, trained model weights were obtained directly from the respective public repositories (ASOptimizer: github.com/Spidercores/ASOptimizer; OligoAI: huggingface.co/barneyhill/OligoAI) without any retraining or fine-tuning.

## SUPPLEMENTAL FIGURES

**Supplementary Figure 1:**
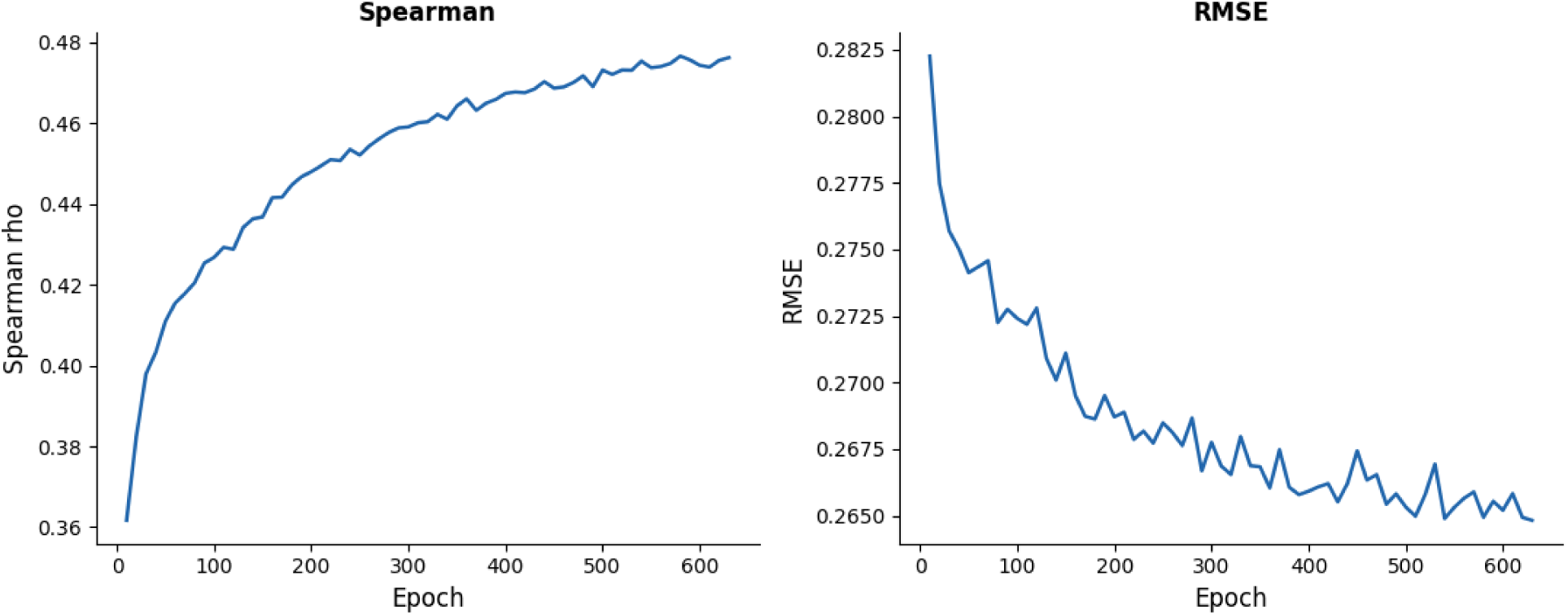
Final model training convergence. The test Spearman rho and root mean squared errors (RMSEs) are shown as a function of the training epoch.

**Supplementary Figure 2:**
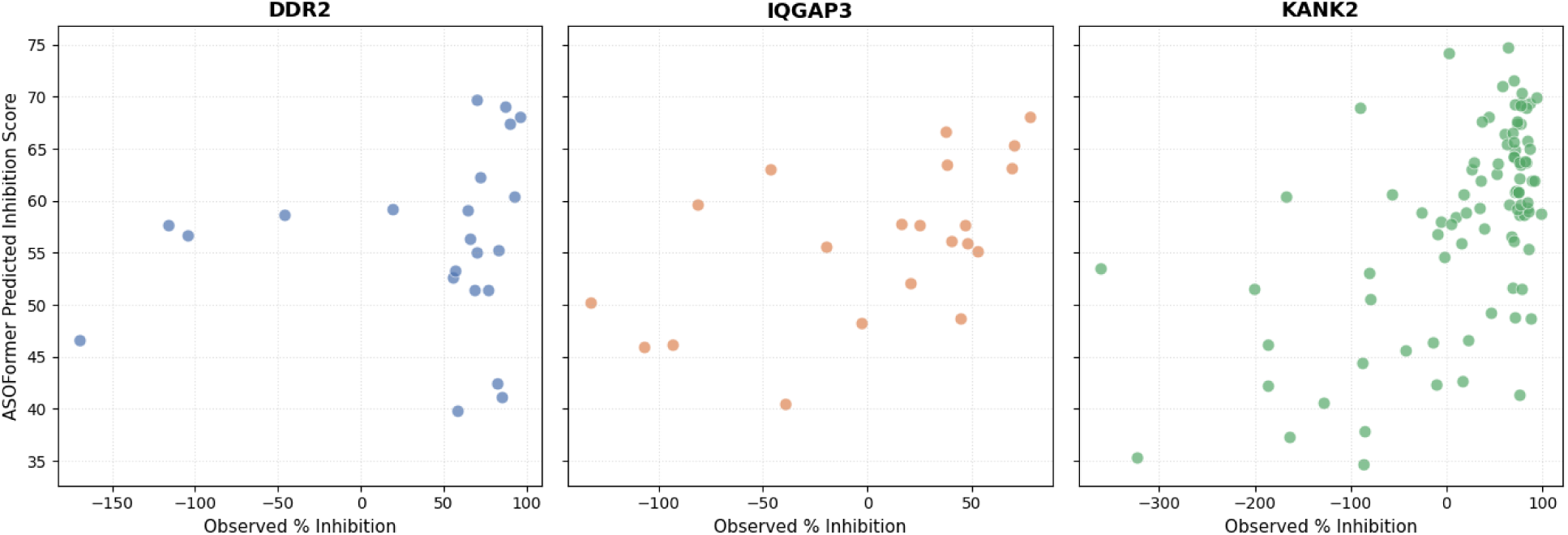
Observed percent inhibition versus ASOFormer predicted inhibition scores for each ASO across the three in-house tested genes.

**Supplementary Figure 3:**
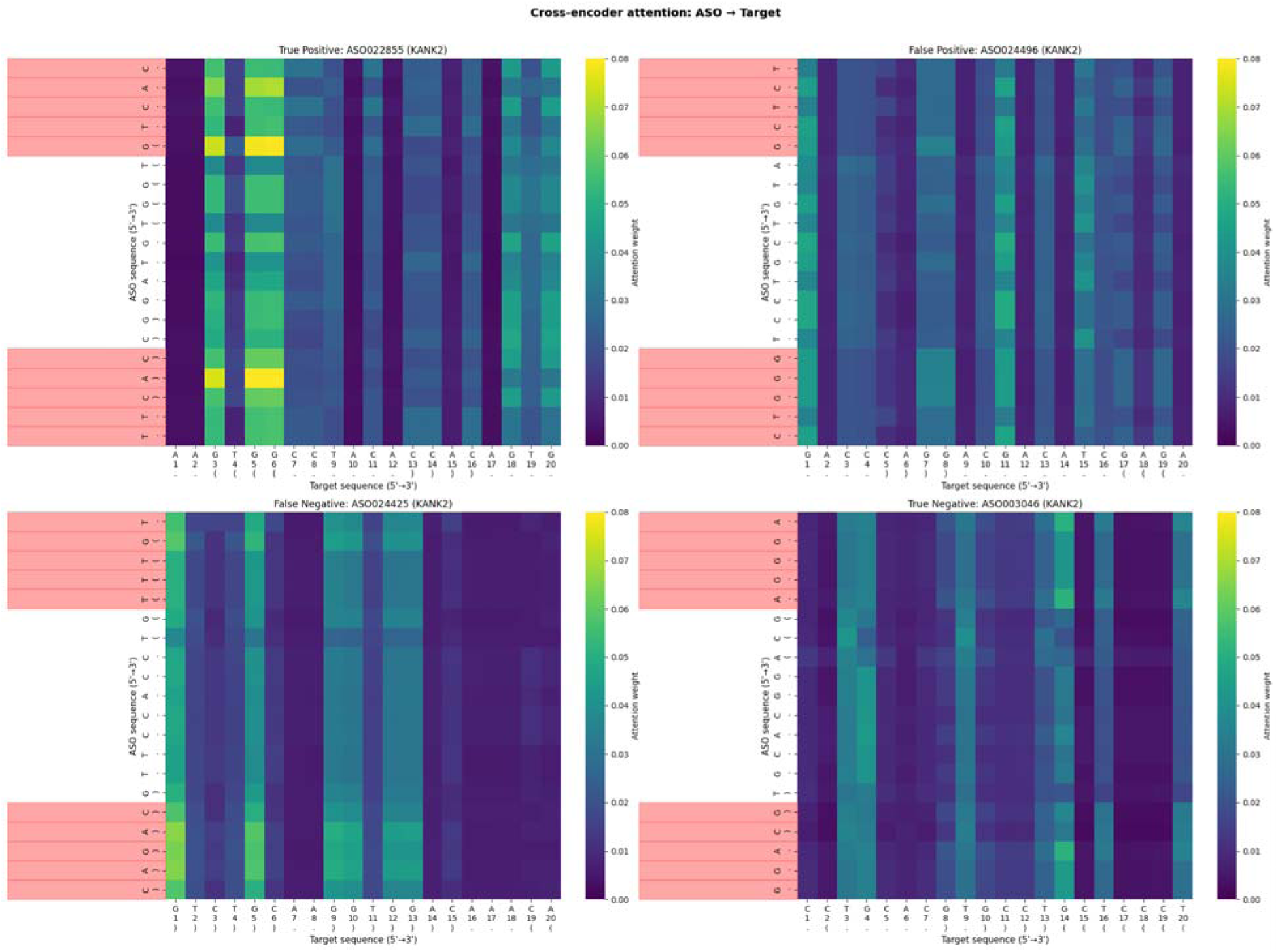
Cross-encoder attention across four *KANK2* ASOs. The four ASOs were chosen to represent different categories of ASOs in the *KANK2* tiling library: true positives (represented in the top 10% of both predicted and observed values), false positives (represented in the top 10% of predicted values but outside of the top 10% of observed values), false negatives (represented in the top 10% of observed values but outside of the top 10% of predicted values), and true negatives (represented in the top 10% of neither predicted nor observed values). Bases overlayed with red on the ASO sequence contain MOE sugar modifications.

**Supplementary Figure 4:**
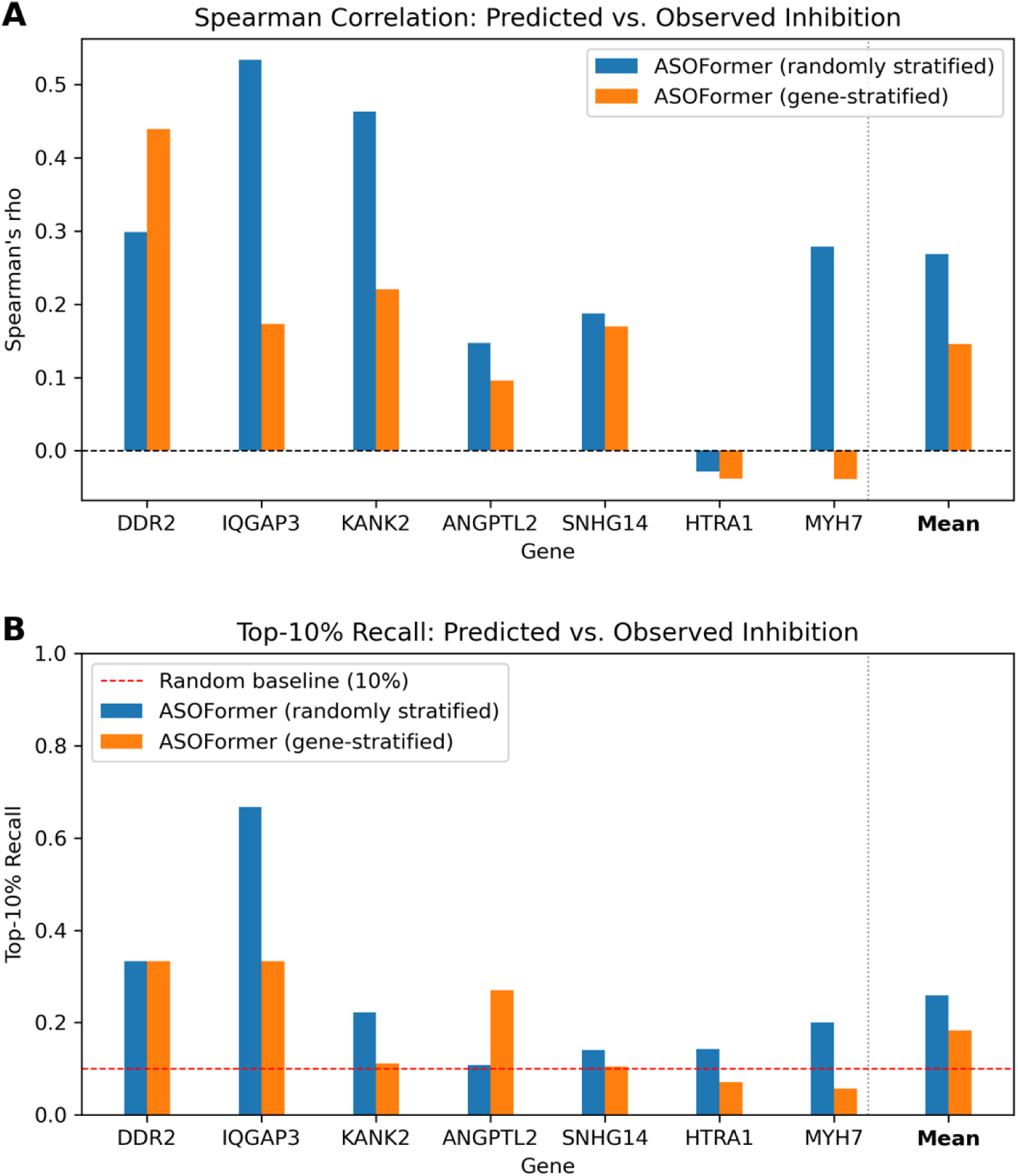
Spearman correlation and top-10% recall of ASOFormer prediction accuracy comparing ASOFormer trained with randomly stratified training/test folds versus ASOFormer trained with gene-stratified training/test folds using knockdown data from seven genes not represented in the training dataset (ASO Atlas).

**Supplementary Figure 5:**
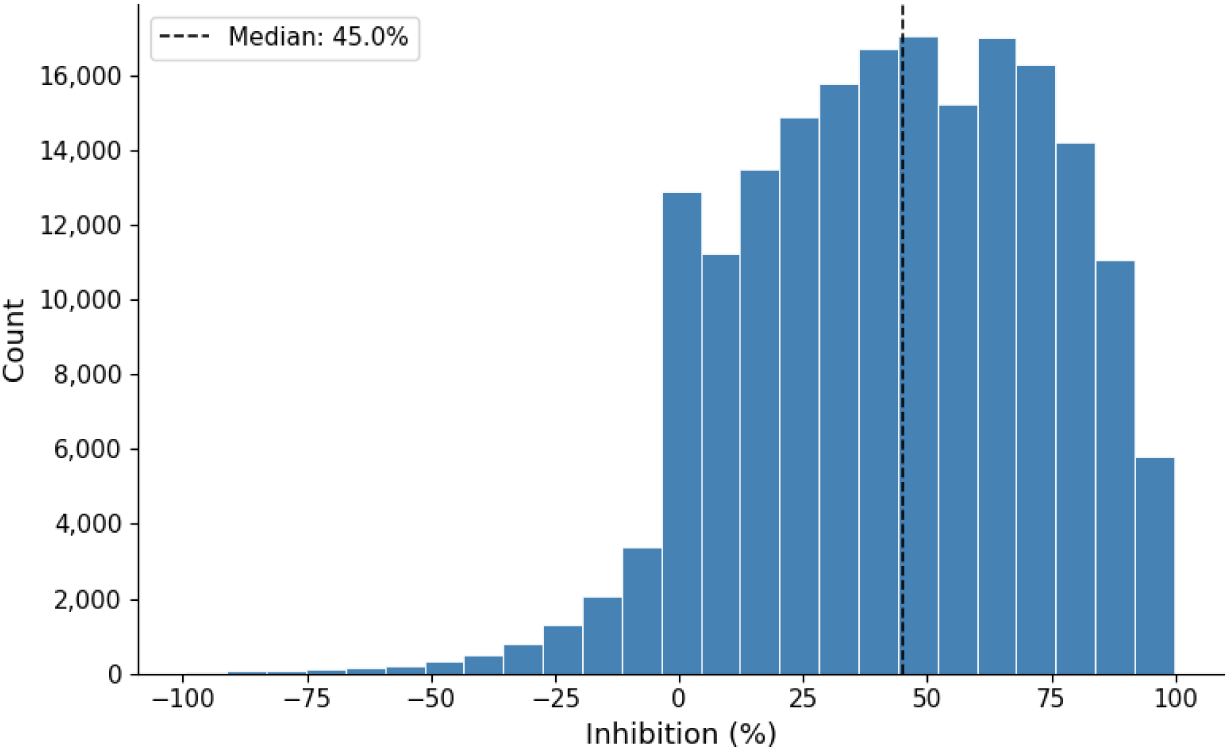
Distribution of ASO inhibition percentages in ASO Atlas. The median (45%) is notated with a dotted line.

## SUPPLEMENTARY TABLES

**Supplementary Table 1:** ASOs designed and tested in-house. Includes observed knockdown values.

<u>ASOFormer_Supp_Tab_1.csv</u>

**Supplementary Table 2:**
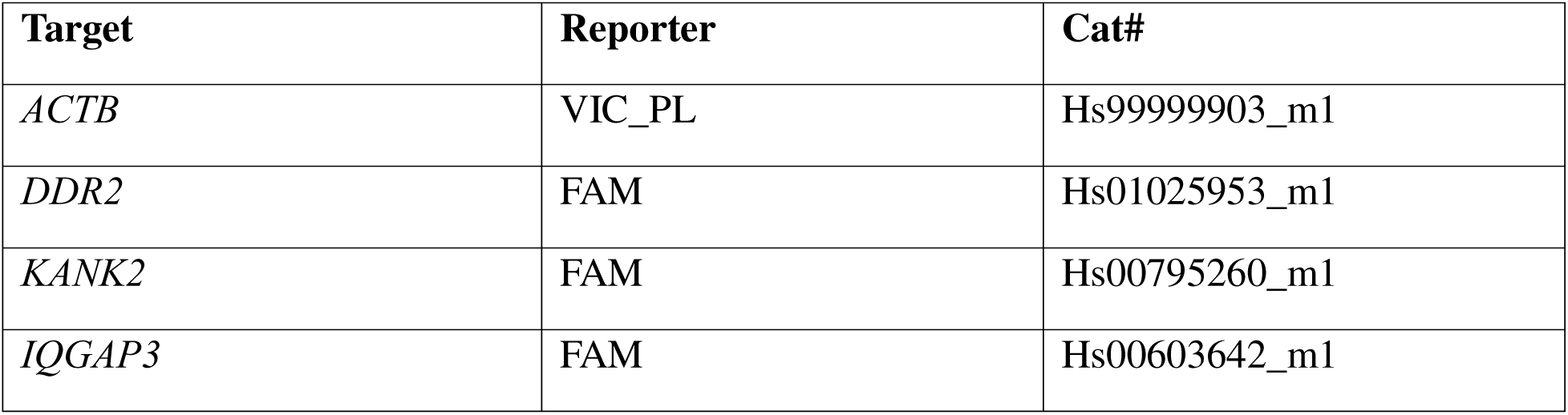
RT-qPCR probes used in this study.

## SUPPLEMENTARY NOTE

### Rationale for random stratification in ASOFormer training

During model development, we considered two approaches to splitting the training and validation data: random stratification, in which ASOs are partitioned without regard to target gene, and gene stratification, in which all ASOs targeting a given gene are held out together. Gene stratification was initially appealing because, in principle, it forces the model to generalize across unseen gene contexts, which might better reflect real-world deployment conditions.

To evaluate this hypothesis empirically, we trained ASOFormer under both conditions and compared performance on a held-out panel of seven external genes. Contrary to our expectation, random stratification yielded consistently superior performance across both evaluation metrics (**Supplementary Figure 4**). The mean Spearman’s rho across the seven genes was 0.27 for random stratification versus 0.14 for gene stratification, and the mean top-10% recall was 0.24 versus 0.15, respectively. Notably, the gene-stratified model produced negative rho values for two of the seven external genes, suggesting that gene stratification may introduce instability or overfitting to gene-level features that do not transfer well.

These findings indicate that, at the current scale of available training data, random stratification provides a better inductive bias for the model and leads to more reliable generalization to unseen genes. Accordingly, all reported ASOFormer models use random stratification.

## REFERENCES

1. Dhuri, K., Bechtold, C., Quijano, E., et al. (2020) Antisense Oligonucleotides: An Emerging Area in Drug Discovery and Development. J Clin Med, 9, 2004.

2. Egli, M. and Manoharan, M. (2023) Chemistry, structure and function of approved oligonucleotide therapeutics. Nucleic Acids Res, 51, 2529–2573.

3. Dowdy, S.F. (2017) Overcoming cellular barriers for RNA therapeutics. Nat Biotechnol, 35, 222–229.

4. Lieberman, J. (2018) Tapping the RNA world for therapeutics. Nat Struct Mol Biol, 25, 357–364.

5. Miller, T.M., Cudkowicz, M.E., Genge, A., et al. (2022) Trial of Antisense Oligonucleotide Tofersen for SOD1 ALS. N Engl J Med, 387, 1099–1110.

6. de Smet, M.D., Meenken, C.J. and van den Horn, G.J. (1999) Fomivirsen - a phosphorothioate oligonucleotide for the treatment of CMV retinitis. Ocul Immunol Inflamm, 7, 189–198.

7. Rader, D.J. and Kastelein, J.J.P. (2014) Lomitapide and mipomersen: two first-in-class drugs for reducing low-density lipoprotein cholesterol in patients with homozygous familial hypercholesterolemia. Circulation, 129, 1022–1032.

8. Finkel, R.S., Mercuri, E., Darras, B.T., et al. (2017) Nusinersen versus Sham Control in Infantile-Onset Spinal Muscular Atrophy. N Engl J Med, 377, 1723–1732.

9. Iuchi, H., Matsutani, T., Yamada, K., et al. (2021) Representation learning applications in biological sequence analysis. Computational and Structural Biotechnology Journal, 19, 3198–3208.

10. Hill, B., Jaques, M.R., Nair, R.R., et al. (2025) Accurately modelling RNase H-mediated antisense oligonucleotide efficacy. 10.1101/2025.10.29.685292.

11. Crooke, S.T., Baker, B.F., Crooke, R.M., et al. (2021) Antisense technology: an overview and prospectus. Nat Rev Drug Discov, 20, 427–453.

12. Bennett, C.F. and Swayze, E.E. (2010) RNA targeting therapeutics: molecular mechanisms of antisense oligonucleotides as a therapeutic platform. Annu Rev Pharmacol Toxicol, 50, 259–293.

13. Rinaldi, C. and Wood, M.J.A. (2018) Antisense oligonucleotides: the next frontier for treatment of neurological disorders. Nat Rev Neurol, 14, 9–21.

14. Hwang, G., Kwon, M., Seo, D., et al. (2024) ASOptimizer: Optimizing antisense oligonucleotides through deep learning for IDO1 gene regulation. Molecular Therapy - Nucleic Acids, 35, 102186.

15. Rotrattanadumrong, R. and Donno, C.D. (2025) OligoGym: Curated Datasets and Benchmarks for Oligonucleotide Drug Discovery. In.

16. Sharad, S. and Kapur, S. (2019) Antisense Therapy BoD – Books on Demand.

17. Chen, E.S. and Ho, E.S. (2023) In-silico study of antisense oligonucleotide antibiotics. PeerJ, 11, e16343.

18. Lorenz, R., Bernhart, S.H., Höner zu Siederdissen, C., et al. (2011) ViennaRNA Package 2.0. Algorithms Mol Biol, 6, 26.

19. McQuisten, K.A. and Peek, A.S. (2007) Identification of sequence motifs significantly associated with antisense activity. BMC Bioinformatics, 8, 184.

20. Matveeva, O.V., Tsodikov, A.D., Giddings, M., et al. (2000) Identification of sequence motifs in oligonucleotides whose presence is correlated with antisense activity. Nucleic Acids Res, 28, 2862–2865.

21. Zhang, L., Vickers, T.A., Sun, H., et al. (2021) Binding of phosphorothioate oligonucleotides with RNase H1 can cause conformational changes in the protein and alter the interactions of RNase H1 with other proteins. Nucleic Acids Res, 49, 2721–2739.

22. Kielpinski, L.J., Hagedorn, P.H., Lindow, M., et al. (2017) RNase H sequence preferences influence antisense oligonucleotide efficiency. Nucleic Acids Res, 45, 12932–12944.

23. Leckie, J. and Yokota, T. (2025) Integrating Machine Learning-Based Approaches into the Design of ASO Therapies. Genes, 16, 185.

24. Hagedorn, P.H., Pontoppidan, M., Bisgaard, T.S., et al. (2018) Identifying and avoiding off-target effects of RNase H-dependent antisense oligonucleotides in mice. Nucleic Acids Res, 46, 5366–5380.

25. Wan, W.B. and Seth, P.P. (2016) The Medicinal Chemistry of Therapeutic Oligonucleotides. J. Med. Chem., 59, 9645–9667.

26. Paszke, A., Gross, S., Massa, F., et al. (2019) PyTorch: An Imperative Style, High-Performance Deep Learning Library. 10.48550/arXiv.1912.01703.

27. Devlin, J., Chang, M.-W., Lee, K., et al. (2019) BERT: Pre-training of Deep Bidirectional Transformers for Language Understanding. 10.48550/arXiv.1810.04805.

28. Hunter, J.D. (2007) Matplotlib: A 2D Graphics Environment. Computing in Science & Engineering, 9, 90–95.

29. Waskom, M.L. (2021) seaborn: statistical data visualization. Journal of Open Source Software, 6, 3021.

